# The tricuspid valve also maladapts: A multiscale study in sheep with biventricular heart failure

**DOI:** 10.1101/2020.09.03.278515

**Authors:** William D Meador, Mrudang Mathur, Gabriella P Sugerman, Marcin Malinowski, Tomasz Jazwiec, Xinmei Wang, Carla MR Lacerda, Tomasz A Timek, Manuel K Rausch

## Abstract

**Objectives:** We set out to determine the tricuspid valve’s propensity to (mal)adapt in disease.

**Background:** Tricuspid regurgitation (TR) is generally considered secondary to right and/or left ventricular disease without organic failure. Interestingly, we and others have previously shown the mitral valve (mal)adapts in functional mitral regurgitation, which may warrant reconsideration of its functional etiology. Whether the tricuspid valve similarly (mal)adapts is mostly unknown.

**Methods:** We evaluated the (mal)adaptive response of tricuspid valve anterior leaflets (TVALs) from an ovine model in which over-pacing (19 ± 6 days) induced biventricular heart failure and TR (tachycardia-induced cardiomyopathy, TIC, n=33) and compared findings to those from a control group (n=17). In both groups, we performed proteomics, immunohistochemistry, histology, two-photon microscopy, collagen assays, leaflet thickness and morphology measurements, and biaxial mechanical tests.

**Results:** We found metabolically active resident valvular cells in TIC TVALs which expressed activation and turnover markers. In TIC TVALs, we observed a 140% increase in collagen content (p=0.016), increased collagen dispersion regionally (p=0.017), a 130% increase in leaflet area (p=0.002), a 140% increase in thickness (p=0.006), and a 130% increase in radial stiffness (p=0.006).

**Conclusions:** Our data suggest that TVALs (mal)adapt during TIC on all scales. This response is likely initiated by activated valvular cells, resulting in collagen turnover, and ultimately leading to thickening, area increase, and stiffening. Our data motivates future studies on the exact pathways leading to tricuspid (mal)adaptation and pharmacological therapeutic strategies for TR.

**Condensed Abstract:** In most cases, tricuspid regurgitation is presumed to originate from valve extrinsic factors. We challenge this paradigm and hypothesize that the tricuspid valve maladapts, rendering the valve at least partially culpable for its dysfunction. As such, we set out to demonstrate that the tricuspid valve, indeed, maladapts in an ovine model of heart disease. In the anterior leaflets, we found alterations on the protein and cell-level, leading to maladaptation in the form of tissue growth, thickening, and stiffening. Our findings may initially motivate mechanistic pathway studies, and in the future, leaflet-targeted pharmacological therapeutic options for tricuspid regurgitation.

## Introduction

Moderate to severe tricuspid regurgitation (TR) affects more than 1.6 million Americans and is an independent predictor of mortality (1–3). In 85% of severe cases, TR is considered functional or secondary to right ventricular (RV) remodeling and left sided heart disease (1,4,5). Here, resultant tricuspid annular dilation and papillary muscle displacement are believed to circumferentially strain and tether the valve’s leaflets, respectively, preventing proper leaflet coaptation (6,7). In other words, it is believed that constraints from valve-extrinsic conditions render the valve dysfunctional, while the valve itself is considered structurally and mechanically intact and healthy (8,9). In fact, TR was historically not treated assuming that it would resolve following treatment of left sided disease (10). However, this conservative approach to TR has since been reconsidered and treatment strategies are more aggressive, including invasive surgical repair or replacement. Despite decades of experience with these devices and techniques, failure rates are still as high as 10-30% (11). As such, TR remains a poorly understood disease with significant opportunity for therapeutic improvement.

Interestingly, on the left side, functional mitral regurgitation has been found to be not so functional after all (12). Our group and others have shown that the mitral valve leaflets grow and remodel in functional mitral regurgitation (13–16). While leaflet growth, i.e., adaptation, in the mitral valve may be beneficial by increasing the coaptation area (17,18), concomitant leaflet fibrosis, i.e., maladaptation, may stiffen the valve and thus impede proper kinematics and coaptation (19). Thus, mitral leaflet (mal)adaptation, i.e., both adaptation and maladaptation, plays a diametrical role. Fortunately, Levine and co-workers have demonstrated that the maladaptive, fibrotic response of the leaflets may be pharmacologically reduced without reducing their adaptive ability to grow (20). Thus, in the future, pharmacological management of mitral regurgitation targeting leaflet remodeling may support its surgical/interventional treatment and improve long-term outcomes (21).

Similar research on the tricuspid valve is nearly absent, so much so that it has been dubbed the “forgotten valve” (22). Although a first study in humans demonstrated that leaflets of patients with pulmonary hypertension may increase in area, it remains unknown whether tricuspid valve leaflets similarly maladapt during functional TR (23). Whether they actively maladapt or not is ultimately important as it may, as is the case with the mitral valve, challenge the notion of TR being strictly functional. In turn, a better understanding of TR may inform improved surgical strategies or provide alternative treatment routes. Therefore, our objective was to fill this gap in knowledge and investigate whether the tricuspid valve anterior leaflet (TVAL), similarly to the mitral valve, (mal)adapts in an ovine, tachycardia-induced cardiomyopathy (TIC) biventricular heart failure model that we have previously used to study valvular disease (24–26).

## Methods

Detailed methods and materials are provided within the online Supplementary Materials.

### Animal model

All aspects of this research study were performed in accordance with the Principles of Laboratory Animal Care, formulated by the National Society for Medical Research, and the Guide for Care and Use of Laboratory Animals prepared by the National Academy of Science and published by the National Institutes of Health. Additionally, this protocol was developed, reviewed, and performed in accordance with the approval of a local Institutional Animal Care and Use Committee.

The TIC ovine model used in this study has been previously described in detail and validated as a reliable and repeatable model of biventricular heart failure in sheep (24). In short, we randomly assigned adult male Dorset sheep to either control (CTL, n=17, 59.9 ± 4.6 kg) or disease (TIC, n=33, 60.1 ± 5.3 kg) groups. In all animals, we monitored baseline (i.e. prior to pacing) ventricular function and valvular competence via epicardial echocardiography. In TIC subjects, we sutured a monopolar pacing lead onto the lateral left ventricular (LV) wall, gaining access through a left lateral mini-thoracotomy approach (10-15 cm, fifth/sixth intercostal space) with a surgical incision approximated in standard fashion, and exteriorized the lead through the thorax to a pacemaker (Consulta CRT-P, Medtronic, Minneapolis, MN). We initiated biventricular failure via a progressive 180-260 beats per minute pacing protocol. Once both moderate TR and LV dysfunction (ejection fraction (EF) < 30%) were present (19 ± 6 days), we performed a terminal (i.e., post-pacing) epicardial echocardiograph. During the terminal procedure of all animals, we acquired hemodynamic data prior to euthanasia. After euthanasia, we isolated the tricuspid valve from each animal to be used in this study. For medications and procedural details, see the Supplementary Materials.

### Morphology and storage

Immediately after explanting the tricuspid valves, we unfolded the valves on a calibrated grid and calculated the TVAL area, major cusp width, and major cusp height from orthogonal photographs. We subsequently stored the tricuspid valves at −80°C (9:1 DMEM:DMSO with protease inhibitor) until rapid thaw to room temperature for testing. Prior to any testing, we removed all chordae tendineae.

### High through-put analysis of protein expression

In brief, we submitted dried peptides from TVAL fragments to the University of Texas at Austin CBRS Biological Mass Spectrometry Facility for protein identification by liquid chromatography tandem mass spectrometry to identify differentially expressed proteins among treatment groups. With these differentially expressed proteins we generated protein interaction maps via STRING v11 (Search Tool for the Retrieval of Interacting Genes/Proteins, STRING Consortium) and identified Gene Ontologies via PANTHER (Protein ANalysis THrough Evolutionary Relationships). For details and previously reported protocols (27,28), see the Supplementary Materials.

### Histology, thickness, and immunohistochemistry

We fixed radial strips of TVALs from annulus to free edge for commercial histological and immunohistochemistry services (HistoServ, Inc., Amaranth, MD). To determine tissue thickness, we used custom MATLAB code to fit splines to the atrialis and ventricularis surface of H&E stained sections and calculated the normal distance between splines. We summarized the measurements into three equidistant regions (i.e. near-annulus, belly, and free edge). Additionally we performed regional analyses on four immunohistochemistry markers previously cited in heart valve growth and remodeling processes: (i) α-smooth muscle actin (αSMA), (ii) Ki67, (iii) matrix metalloproteinase 13 (MMP13), and (iv) transforming growth factor β1 (TGF-β1) (14,15,29,30). Furthermore, we were specifically interested in the location of these markers as regions of tissue may be differentially active, due to regionally varying stimuli and composition (31–33). For regional analyses, we interpolated 10 thickness regions between the atrialis and ventricularis splines in all three length regions (i.e. 30 total regions), quantifying positively stained pixels using a custom MATLAB program. For details, see the Supplementary Materials.

### Quantitative collagen assay

We acquired the wet mass of tissue samples from annulus, belly, and free edge regions of TVALs. We then quantified wet mass collagen content using a total collagen assay kit (Biovision Inc., K406, Milpitas, CA, USA). For details, see the Supplementary Materials.

### Biaxial testing and analysis

Using a commercial biaxial testing device (Biotester, Cellscale, Waterloo, ON, Canada), we mechanically tested 7mm × 7mm square samples from the belly region of TVALs as previously described (31). We characterized nonlinear mechanical stretch-membrane tension curves with four metrics: (i) toe stiffness (i.e., lower region slope), (ii) calf stiffness (i.e., slope near 20 N/m) (iii) heel stretch, (i.e., stretch where the toe region transitions into the calf region), and (iv) anisotropy index, (i.e., ratio between the circumferential and radial stretches at 20 N/m). For details, see the Supplementary Materials.

### Two-photon microscopy

We analyzed the collagen microstructure and cell nuclei morphology using two-photon microscopy methods previously described (31,34). For details, see the Expanded Materials and Methods in the Supplementary Materials.

### Statistical analyses

For all experimental data, we first performed Shapiro-Wilk tests to determine whether our data were normally distributed. Additionally, we tested whether the variances of our data sets were similar through an F-test. Under all disease versus control comparisons, a Student’s t-test was used if the data were normally distributed and had similar variances. Otherwise, we used the Wilcoxon Rank Sum Test or Welch’s t-test, as appropriate. In correlations, if either variable failed the normality assumption or was ordinal in type, we used the nonparametric Spearman rank correlation. Otherwise, we used a Pearson correlation. For statistical comparisons and correlations we used either one-tailed or two-tailed tests, as appropriate, with our stated hypotheses. We defined a p-value less than 0.05 as significant. All statistical comparisons and correlations were implemented in MATLAB.

## Results

### Animal model outcomes

Of the 17 CTL animals, we successfully acquired echocardiographic data from 14 animals and hemodynamic data from 13 animals. Of the 33 TIC animals whose tissue we used in this study, we successfully acquired echocardiographic data from 28 animals and hemodynamic data from 30 animals. Of the 28 animals with echocardiographic data, 12 developed both moderate or greater TR and < 30% LV EF (i.e., our end-pacing criteria for the model), 11 only developed moderate or greater TR (without < 30% LV EF), 2 only developed < 30% LV EF (without moderate or greater TR), and 3 did not develop either. No TIC subjects were excluded.

Refer to Table 1 for echocardiographic data of CTL and TIC (baseline and terminal) groups and hemodynamic data of CTL and TIC (terminal) groups. After pacing, echocardiography revealed significant parameter changes consistent with clinical biventricular dysfunction and functional TR. To summarize, we observed the TIC (terminal) group to have a significant reduction in RV EF, LV EF, and RV fractional area change, and significantly increased TR grade, mitral regurgitation grade, RV inner dimension at diastole (IDd), LV IDd, and tricuspid valve annulus dimension. Hemodynamically, TIC animals had significantly increased heart rates and RV maximal and end-systolic pressures.

**Table 1:** Echocardiographic and hemodynamic data of animal model

| Parameter | CTL | TIC (baseline) | TIC (terminal) |
| --- | --- | --- | --- |
| *Echocardiographic data* |  |  |  |
| TR grade | 1.0 (0.0) | 0.0 (0.0) | 2.0 (1.0) * † |
| RV EF, % | 67.0 ± 8.6 | 58.8 (20.5) | 48.4 ± 12.9 * † |
| RV FAC, % | 52.1 ± 6.0 | 53.9 ± 8.0 | 37.4 ± 8.7 * † |
| RV IDd, cm | 2.5 ± 0.4 | 2.6 (0.6) | 2.9 ± 0.7 * |
| TV annulus dimension, cm | 2.5 (0.3) | 2.5 ± 0.4 | 3.2 ± 0.5 * † |
| MR grade | 0.0 (0.0) | 0.0 (0.0) | 2.0 (1.0) * † |
| LV EF, % | 55.8 ± 4.4 | 61.3 ± 6.2 | 30.0 (11.2) * † |
| LV IDd, cm | 4.2 ± 0.3 | 3.7 ± 0.6 | 4.6 ± 0.4 * † |
| *Hemodynamic data* |  |  |  |
| HR, bpm | 108 (11) | - | 126 ± 23 * |
| RV P<sub>Max</sub>, mmHg | 28.8 (11.4) | - | 44.0 ± 11.5 * |
| RV P<sub>ES</sub>, mmHg | 22.3 ± 5.6 | - | 35.7 ± 11.0 * |
| RA P<sub>Mean</sub>, mmHg | 10.5 (3.2) | - | 10.6 (5.6) |
| LV P<sub>Max</sub>, mmHg | 96.1 ± 9.7 | - | 92.9 ± 13.8 |
| LV P<sub>ES</sub>, mmHg | 63.1 ± 17.8 | - | 64.4 ± 13.6 |

Values are mean ± standard deviation or median (interquartile range).
CTL=control, EF=ejection fraction, ES=end systolic, FAC=fractional area change, HR=heart rate, IDd=inner dimension at diastole, LV=left ventricular, Max=maximal, MR=mitral regurgitation, P=pressure, RA=right atrial, RV=right ventricular, TIC=tachycardia-induced cardiomyopathy, TR=tricuspid regurgitation, TV=tricuspid valve.
\* p < 0.05 vs. CTL, † p < 0.05 vs TIC (baseline)

### TVAL area increases in TIC

Once isolated, all TVALs appeared anatomically typical (Figure 1a) (35). However, in our analysis we found a significant increase of approximately 130% in TVAL area for TIC animals compared to CTL animals (p=0.002, Figure 1b). These increases in area appeared to be driven by a significant increase in width (p=0.023), and a near significant increase in height (p=0.070) (Figure 1c-d). Furthermore, the TVAL areas showed no correlation with animal weight in either animal groups (CTL: r=0.239, p=0.253, TIC: r=0.077, p=0.349) (Supplementary Figure 1). In summary, TIC TVALs areas were 130% larger when compared to CTL TVALs, ostensibly driven by an increase in leaflet width.

**Figure 1:**
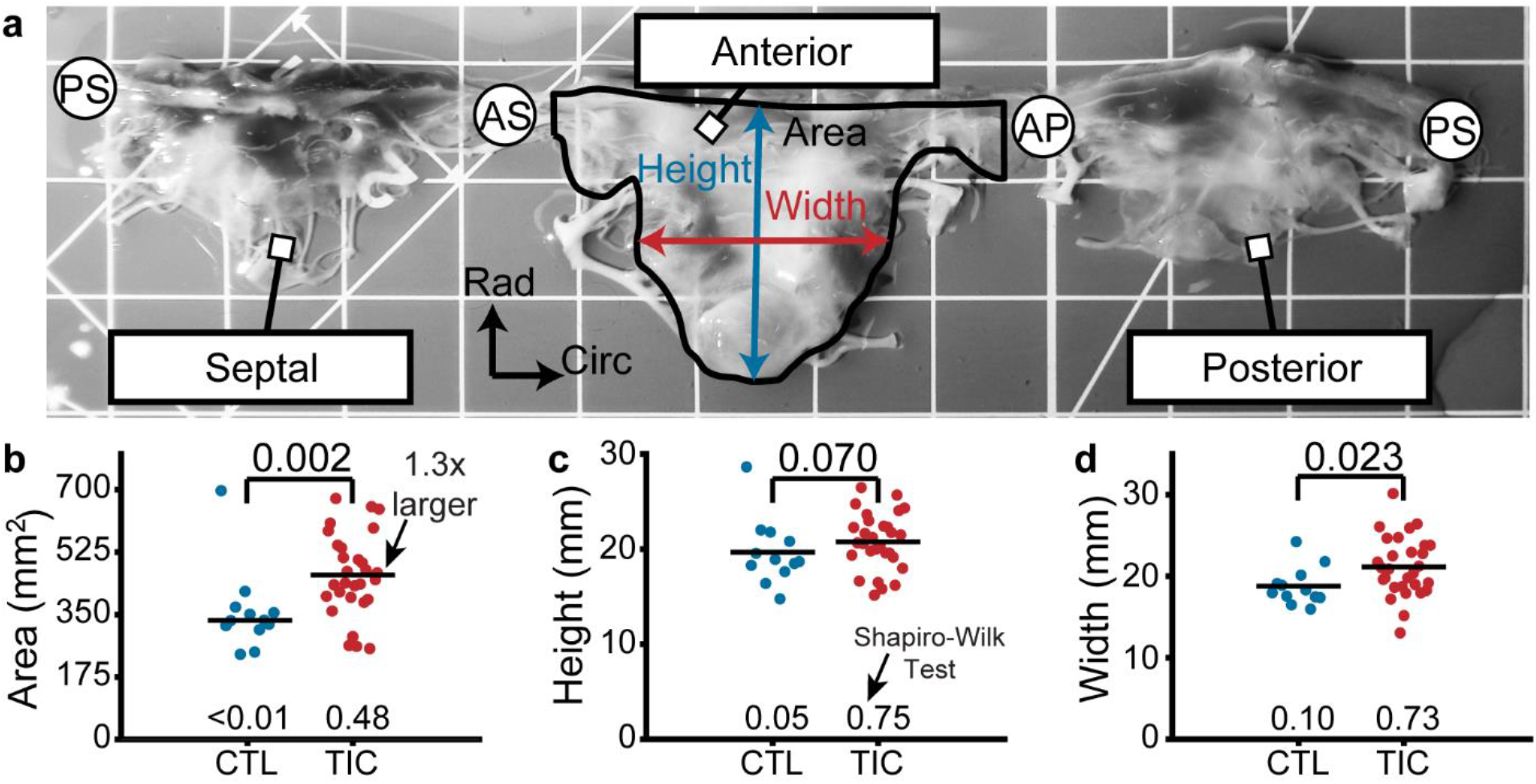
Anterior leaflet area increases in tachycardia-induced cardiomyopathy (TIC). **(a)** An ovine tricuspid valve unfolded at the postero-septal commissure (PS). Located between the antero-septal (AS) and antero-posterior (AP) commissures, the anterior leaflet and its measured area (black), height (blue), and width (red) are shown. Grid scale=1 cm. **(b-d)** Comparisons between control (CTL, blue) and TIC (red) anterior leaflet **(b)** area, **(c)** height, and **(d)** width. Black bars represent data mean if normal, and data median if non-normal as determined by Shapiro-Wilk test p-value below data. Values above data represent p-values from Student’s t-test or Wilcoxon Rank Sum test, as appropriate

### TVALs thicken with TIC

We measured TVAL thickness from fixed radial strips of tissue (Figure 2a). In TIC and CTL animals the mean thickness of all TVALs decreased from the near-annulus region (thickest) to the free edge (thinnest) (Figure 2b). Furthermore, the mean thickness of TIC TVALs was consistently larger than CTL TVALs in all three regions. When summarized into a single average thickness, we found a significant increase (140%) in TIC TVAL thickness (p=0.006, Figure 2c). When analyzed by region, the increases in thickness were most strongly driven by a significant increase in the free edge region (p=0.003, Figure 2d). In summary, TIC TVALs were 140% thicker when compared to CTL TVALs, mostly driven by increases in the free edge thickness.

**Figure 2:**
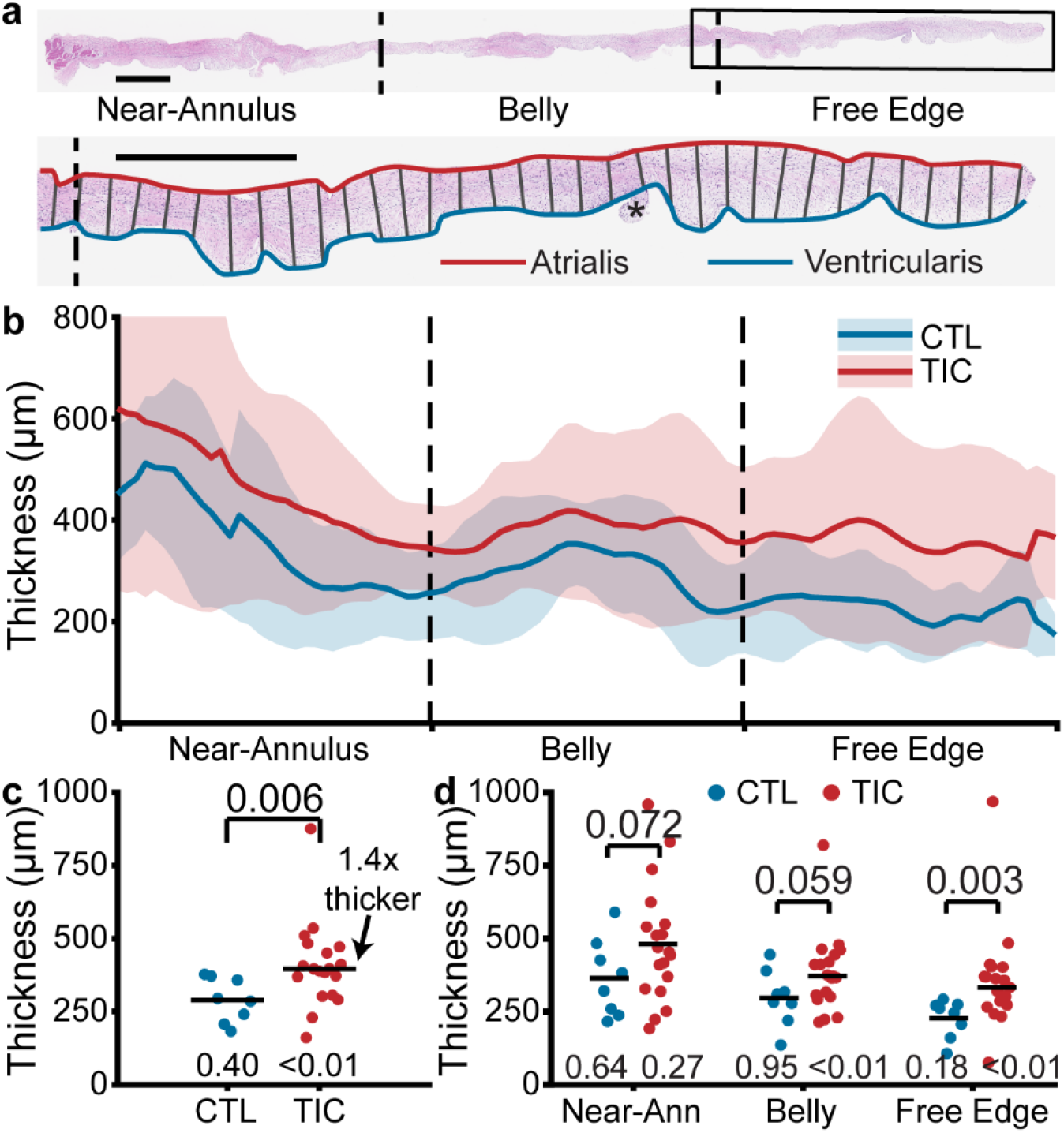
Anterior leaflet thickness increases in tachycardia-induced cardiomyopathy (TIC). **(a)** (top) Representative fixed radial tissue strip approximated into regions of near-annulus, belly, and free edge. Inscribed box (black) is magnified (below) to show thickness measurement between atrialis (red) and ventricularis (blue) splines. Chordae tendineae (*) were excluded manually. Scale bars=1 mm. **(b)** Profiles of control (CTL, blue) and TIC (red) anterior leaflet thickness approximated into equal-third regions of near-annulus, belly, and free-edge. Pictured are mean (solid) +/− 1 standard deviation (shaded). **(c-d)** Thickness comparisons between CTL and TIC groups when data is **(c)** pooled across all regions and **(d)** pooled within regions. Black bars represent data mean if normal, and data median if non-normal as determined by Shapiro-Wilk’s test p-value below data. Values above data represent p-values from Student’s t-test or Wilcoxon Rank Sum test, as appropriate

### Increased metabolic and regulatory proteins in TIC TVALs

Using proteomics, we successfully identified 247 differentially expressed proteins between CTL and TIC TVALs (Supplementary Figure 2). Refer to Supplementary Table 1 for a complete list of FASTA headers, gene names, family information, and expression levels for each of the 247 proteins identified. To better identify the molecular functions of the significant proteins, we built an interactome for easier visualization and interpretation of protein cluster interactions (Supplementary Figure 3). Compared to CTL, TIC TVALs overexpressed metabolic proteins (ENO1, ESD, ALDOA, ALDOC, ASPH, UGP2, AK1, PGK1, PGK2, PKM, LDHA, PGD, among others), serpins (A1, A5, C1, D1, and others), apolipoproteins (A1 and B), and complement proteins (C2, C5, C6 and other subunits). Conversely, underexpressed proteins included proteoglycans (HAPLN1, HAPLN3, ACAN, LUM, KERA and others), ribosomal proteins (S8, S13, S14, S19 and others) and G protein subunits (αi1, αi2, αi3, 11, among others). We further classified the 247 differentially expressed proteins according to their primary families and gene ontologies (i.e. protein classes, molecular functions, cellular component, and biological process). These 247 differentially expressed proteins most often belonged to protein classes of metabolite interconversion enzymes, protein binding modulators, and protein modifying enzymes. Furthermore, these 247 proteins were most associated with the molecular functions of catalytic, binding, and regulatory activity. The 247 proteins’ most common cell component were organelles, protein complexes or membranes. Finally, the most common biological processes for these 247 proteins were metabolic, biological regulation or biogenesis (e.g., histones, ribosomal and membrane proteins). Refer to Supplementary Table 1 for all protein family identifiers, and protein family and gene ontology classifications. In summary, we found 247 differentially expressed proteins in TIC TVALs when compared to CTL TVALs, including many proteins suggestive of increased metabolic and regulatory processes.

### Expression of remodeling associated cellular markers in TIC

We investigated the presence of four cellular markers for remodeling via immunohistochemistry: (i) αSMA, (ii) Ki67, (iii) MMP13, and (iv) TGF-β1 (14,15,29,30). We found an increase in TIC αSMA expression, indicative of cellular activation, mostly in the atrialis of near-annulus and belly regions (Figure 3a). Furthermore, Ki67, a marker for cell proliferation, was increased for TIC in the atrialis of the belly region and in much of the free edge region (Figure 3b). However, we observed only marginal changes in TIC cell nuclei density in the near-annulus (increased) and free edge (decreased) regions (Supplementary Figure 4). Additionally, MMP13, a collagenase, was widely increased for TIC in both near-annulus and belly regions (Figure 3c). Lastly, TGF-β1 also increased primarily in the belly region of TIC samples (Figure 3d). It should be noted that CTL tissues also expressed these markers, albeit to a lesser degree. Overall, we observed regional increases in cellular markers associated with tissue remodeling processes.

**Figure 3:**
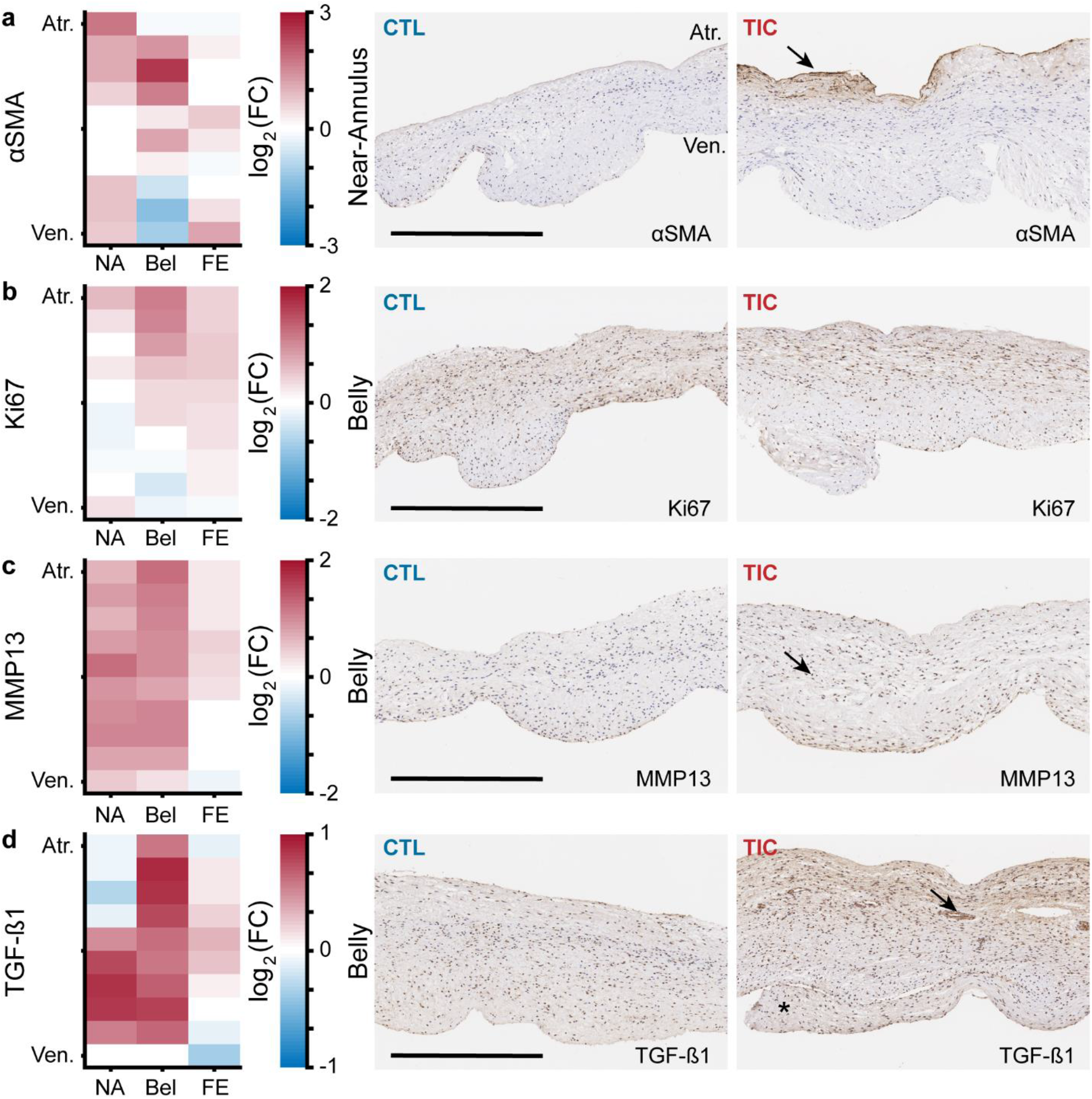
Remodeling cellular markers present in tarchycardia-induced cardiomyopathy (TIC) leaflets. **(a-d)** Heat maps (left) showing regional expression of **(a)** αSMA, **(b)** Ki67, **(c)** MMP13, **(d)** TGF-β1. Heat maps are separated by regions in radial (near-annulus (NA), belly (Bel), and free edge (FE) and thickness (atrialis (Atr.), and ventricularis (Ven.) axes. Logarithm base 2 of the fold change (FC) between control (CTL, n=6) and TIC (n=6) can be interpreted as (positive, red): TIC expression is higher than CTL, (0, white): TIC and CTL expression are approximately equal, and (negative, blue): TIC expression is less than CTL. Representative images of CTL (middle) and TIC (right) are shown with atrialis surface upward. Black arrows indicate **(a)** increased TIC αSMA expression near the atrialis, **(c)** increase in TIC positively stained nuclei for MMP13, and **(d)** interstitial collection of TGF-β1 in potential micro-vessels. Asterisk (*) denotes chordae tendineae excluded from analysis. Scale bars=500 μm

### Collagen content increases in the TVAL in TIC

We quantified the wet tissue collagen content in the near-annulus, belly, and free edge regions of tissue. In both groups, we found collagen content decreased from the near-annulus region to the free edge (Figure 4a). Importantly, in all three regions, the mean collagen content was larger in TIC subjects than in CTL subjects. The strongest increase was in the free edge region (~150%, p=0.006) and the near-annulus region (~135%, p=0.023), but we failed to find a significant increase in the belly region (~125%, p=0.058). When we pooled all regions within subjects, we found a significant 140% increase in collagen content in TIC TVALs (p=0.016, Figure 4b). We also found a strong positive (r=0.650) and significant (p=0.015) correlation between collagen content and TR severity (Supplementary Figure 5). In summary, TIC TVALs had 140% more collagen than CTL TVALs, mostly driven by increases in the free edge and near-annulus content.

**Figure 4:**
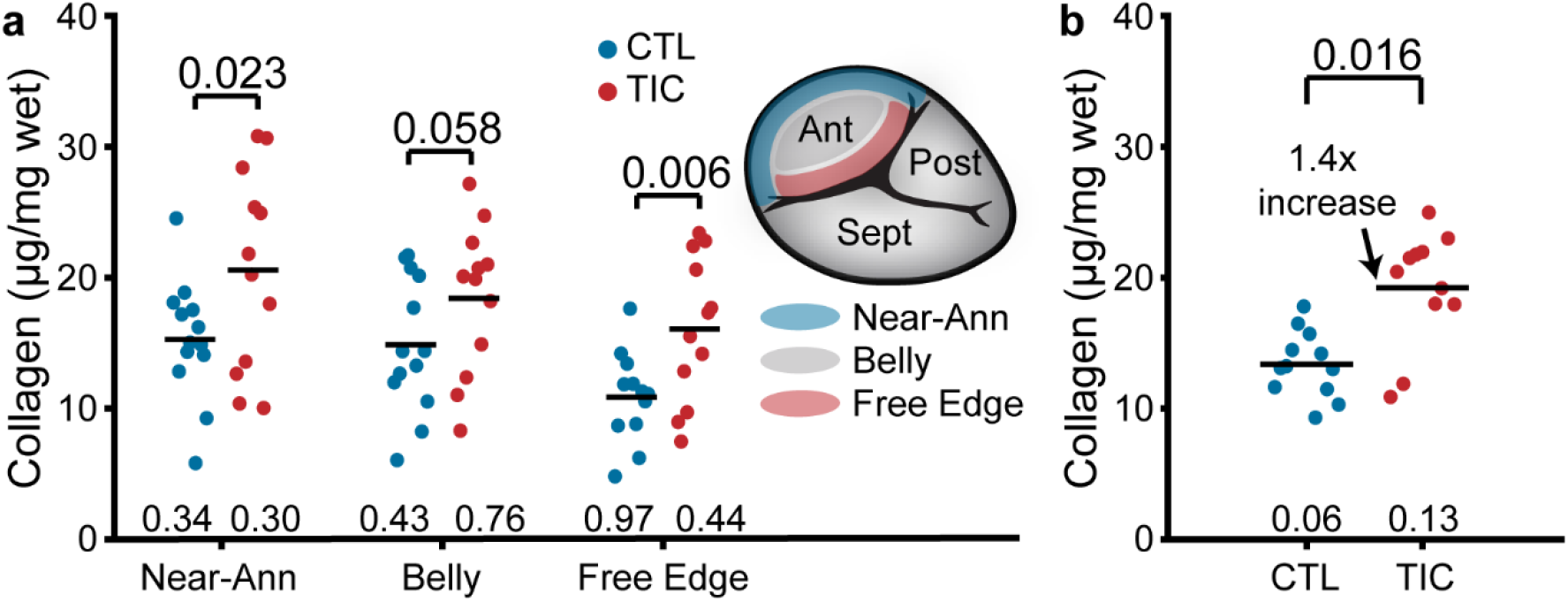
Increased collagen content in tachycardia-induced cardiomyopathy (TIC) anterior leaflets. **(a)** Wet weight collagen content from quantitative collagen assay comparisons between control (CTL, blue) and TIC (red) groups with tissue samples from near-annulus, belly, and free edge regions with inscribed region visualization on tricuspid valve anterior leaflet, and **(b)** when regions are averaged across subjects. Black bars represent data mean if normal, and data median if non-normal as determined by Shapiro-Wilk test p-value below data. Values above data represent p-values from Student’s t-test

### TVALs stiffen in TIC

Under biaxial testing, all tissues exhibited classic J-shaped loading curves, consistent with other biological collagenous tissues (Figure 5a) (36). In detail, the transition stretches and degrees of anisotropy of the curves and remained mostly consistent between CTL and TIC subjects, overall behaving stiffer in the circumferential direction than in the radial direction (i.e., degree of anisotropy < 1) (Supplementary Figure 6). However, our major finding from mechanical characterization was a significant 130% (p=0.006) increase in TIC TVAL stiffness at large stretches (calf stiffness) of the radial loading curves, while circumferential calf stiffness decreased (opposite our hypothesis) (Figure 5b). Additionally, we found that TIC TVALs had significantly increased circumferential stiffnesses at small strains (toe stiffness) (p=0.008, Figure 5c). In summary, TIC TVALs increased in stiffness. They increased predominantly in radial directions at higher stretches, and circumferential directions at lower stretches.

**Figure 5:**
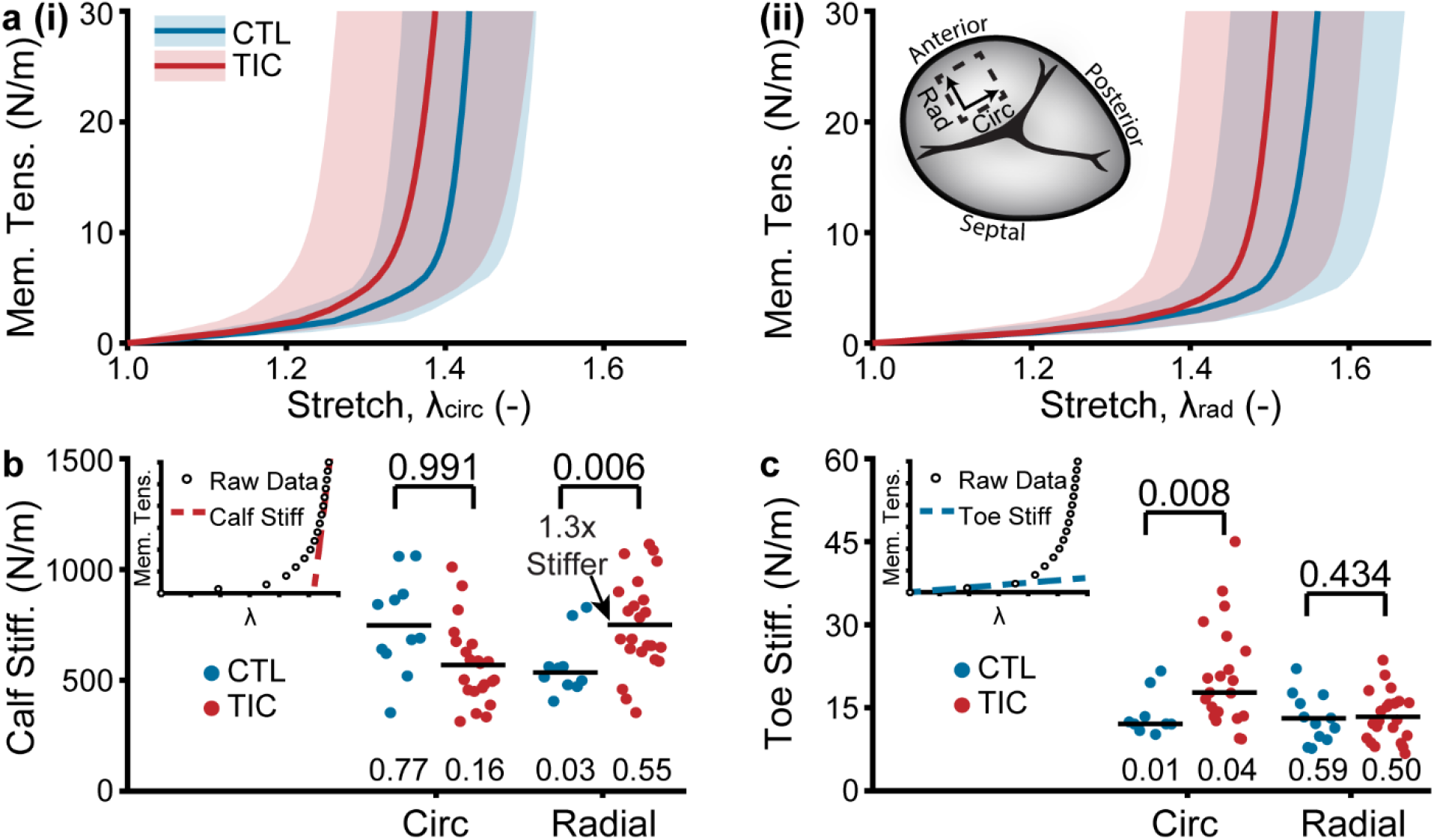
Tachycardia-induced cardiomyopathy (TIC) anterior leaflets are stiffer in radial direction. **(a)** Control (CTL, blue) and TIC (red) membrane tension (Mem. Tens.) vs. stretch average curves (solid) with standard deviation (shaded) in **(i)** circumferential and **(ii)** radial directions. Inset in **(a)** is a visualization of where biaxial samples were acquired (dotted line) from anterior leaflets. **(b-c)** Comparisons of the **(b)** stiffness at large stretches (calf stiffness) and **(c)** stiffness at small stretches (toe stiffness) in circumferential and radial directions. Inset in **(b)** and **(c)** is the definition of calf stiffness (red, dashed) and toe (blue, dashed) we use to characterize a nonlinear material stiffness. Black bars represent data mean if normal and data median if non-normal as determined by Shapiro-Wilk’s test p-value below data. Values above data represent p-values from Student’s t-test or Wilcoxon Rank Sum test, as appropriate

### Altered collagen fiber dispersion in the TIC TVAL atrialis

Utilizing two-photon microscopy, we visualized the collagen and cell nuclei distributions and orientations throughout the entire TVAL thickness. We found that throughout the tissue depth, the mean collagen orientation remained mostly circumferential in CTL and TIC animals, with no apparent changes in mean direction among groups (Figure 6a). However, we noted an increase in collagen fiber dispersion between CTL and TIC (indicated by a larger concentration parameter *κ*). In other words, in TIC animals, more fibers were oriented in the radial direction than in the CTL subjects. When separating our analysis into three depth regions we found a statistically significant increase in fiber dispersion in the 0-33% depth, nearest the atrialis, (p=0.017) in TIC subjects (Figure 6b). Similar analyses in cell nuclei for nuclear orientation, nuclear aspect ratio (NAR), and circularity showed no notable differences between CTL and TIC cell nuclei (Supplementary Figure 7). In summary, we observed that collagen fibers in TIC TVALs were more radially dispersed than CTL TVALs near the atrialis.

**Figure 6:**
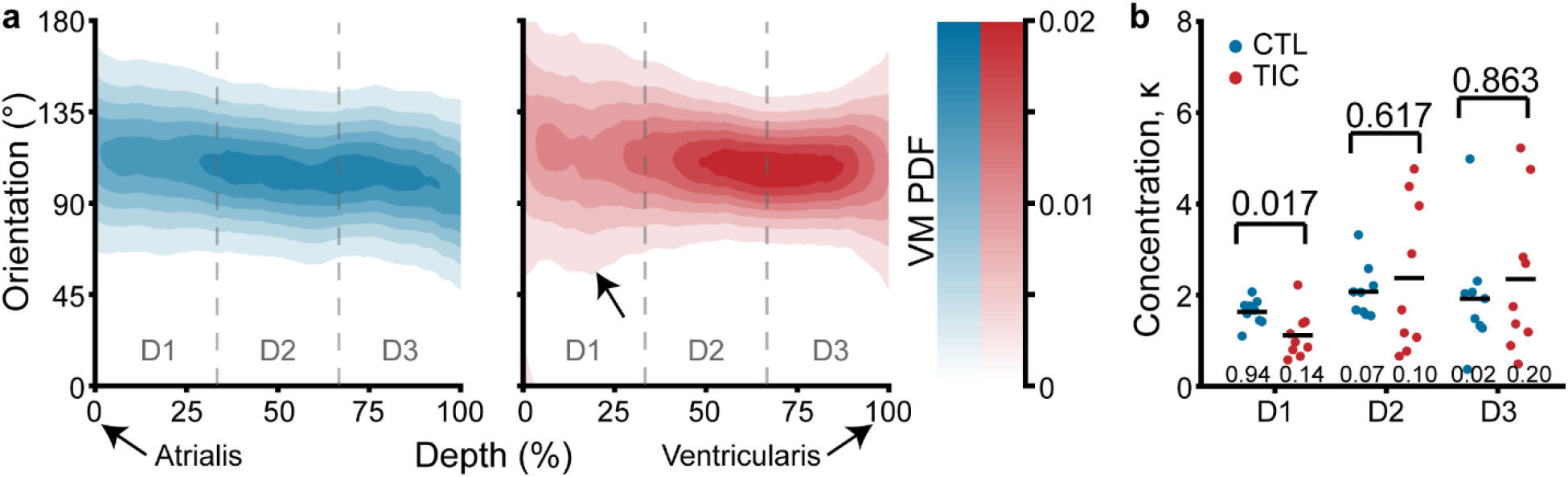
Through-depth collagen microstructure reveals similar mean orientations and regional concentration changes throughout depth between control (CTL) and tachycardia-induced cardiomyopathy (TIC). **(a)** Heat map visualizations of von Mises probability distribution functions (VM PDF) fit to the collagen fiber orientation histograms of two-photon acquired images throughout the entire depth (0% - Atrialis surface, 100% - Ventricularis surface) of averaged CTL (blue) and TIC (red) tissue samples. Orientations of 90° align circumferentially, while 0°/180° align radially. We observed qualitative regional concentration (i.e., heat map width) differences (arrow), which we quantified by **(b)** averaging VM concentration parameter, κ, across depth regions D1, D2, and D3, by subject. Black bars represent data mean if normal, and data median if non-normal as determined by Shapiro-Wilk test p-value below data. Values above data represent p-values from Student’s t-test, Welch’s t-test or Wilcoxon Rank Sum, as appropriate

## Discussion

We investigated the tricuspid valve’s propensity to (mal)adapt to TR, which was motivated by research on mitral valve (mal)adaptation to physiological (37) and pathological stimuli (29,38–40). In the latter setting, we and others have shown that functional mitral regurgitation is not so functional after all (12–16). To date, we have a preliminary, theoretical framework to link mitral leaflet (mal)adaptation to functional mitral regurgitation on multiple scales. We think that pathological mitral leaflet strain initiates a (mal)adaptive cascade with valvular interstitial cell activation and endothelial cell transdifferentiation (38,41,42). These cell populations subsequently increase collagen turnover, and lead to tissue thickening and stiffening (14,15,19). In turn, these tissue changes may contribute to coaptation incompetence. Our current study sought to determine whether the tricuspid valve, too, (mal)adapts on all functional scales. We present our findings in four scales: protein, cell, matrix, and tissue.

On the protein scale, we have identified 247 differentially expressed proteins in TIC TVALs, which we matched to several protein clusters. The increased metabolic protein expressions we observed may suggest increased energetic demands from valvular cells to TIC. This energy may be necessary for the synthesis of apolipoproteins, serpins and complement proteins, which increased in expression. Increases in expression levels of such proteins have been observed in calcific (43) and stenotic (44,45) aortic valves. Such increases may lead to imbalances in lipid metabolism, increased inhibition of proteases such as antithrombins (46,47) with consequent regulation of complement activation (48,49). Overall, the proteomic data indicate that TVAL cells actively respond to TIC.

On the cell scale, using immunohistochemistry we have shown that expression of markers αSMA, Ki67, MMP13 and TGF-β1 were increased in TIC. As a marker for valvular interstitial cell (VIC) activation, increased αSMA suggests a fibrotic response and matrix remodeling (50), consistent with our findings of increased collagen content. Conjunctively, increased MMP13 highlights extracellular turn-over activity through collagenases. While an increased Ki67 expression indicates cellular proliferation, further cellular recruitment from endothelial-to-mesenchymal transition (EndoMT) processes is likely with increased TGF-β1, as TGF-β is a mediator of EndoMT (51) and VIC activation (52) in heart valves. Despite evidence of cellular proliferation and recruitment, cellular density of our tissues remained mostly similar, perhaps due to a balance by the increase in leaflet volume (i.e., area and thickness increase). Furthermore, these cell marker changes were regionally specific suggesting a heterogeneous cellular response, perhaps due to varying regional stimuli.

On the matrix scale, we found strong evidence of increased collagen content as well as an increase in collagen dispersion, i.e., disorganization, near the atrialis surface. The collagen increase may be attributed to the increased fibrotic activity on the cellular scale. Regionally, we observed that collagen increases were most prominent in the near-annulus and free edge regions. Considering the proximity of the near-annulus leaflet tissue to the annulus and the abundance of chordal attachments near the leaflet free edge, we propose that annular dilation and chordal tethering resulting from ventricular remodeling mechanically deformed these regions of leaflet beyond normal physiological levels, inciting an increased profibrotic response in an attempt to maintain mechanobiological homeostasis (53). Secondarily, we observed a more radially dispersed collagen orientation in the atrialis region of the leaflets. Although this may be evidence of radial collagen deposition, with subsequent radial stiffening due to increased radial leaflet stresses from chordal tethering, it is also possible that altered shear stresses from regurgitant blood flow across the atrialis surface induced this microstructural adaptation.

At the tissue scale, we observed leaflet area increase, thickening, and calf region stiffening in the radial direction. Importantly, as opposed to in vivo measurements of area increase that are taken under mechanical stress, our measurements were taken in a mechanically stress-free state. Thus, area increase is entirely due to growth as opposed to elastic stretch. Additionally, we found that tissue calf region stiffness increased in the radial direction and seemingly reduced in the circumferential direction. These changes may be direct responses to annular dilation and papillary muscle displacement in TIC animals that respectively alter circumferential and radial strain. In turn, altered strain may disrupt the mechanobiological equilibrium in those directions, eliciting increased collagen deposition (radial) or degradation (circumferential) in an attempt to reestablish homeostasis (53). Our stiffness results agree with our microstructural observations of increased collagen fiber dispersion in the radial direction and may suggest radial stiffening is an adaptive mechanism of the TVAL. Interestingly, we did observe a significantly stiffer circumferential toe region, which may suggest a role in remodeling for elastin. Additionally, tissue thickness increase may be a direct consequence of upregulated collagen deposition but inflammation-induced tissue swelling cannot be ruled out.

These findings are clinically significant. Similar findings to ours in the mitral valve have inspired the study of pharmacological treatment strategies to reduce leaflet maladaptation, i.e., detrimental tissue effects, while maintaining its adaptive, i.e., beneficial, growth. Specifically, the use of Losartan - an angiotensin II type 1 receptor antagonist that indirectly blocks the TGF-β1 phosphorylation of ERK in EndoMT (51) - resulted in reduced mitral valve leaflet thickness, EndoMT, cellular activation, cell recruitment, and collagen deposition, while preserving leaflet area growth (20). We hope that we have taken a first step toward similar studies that may support pharmacological support of TR surgery and/or intervention. Interestingly, recent work has demonstrated that not only disease, but also surgical repair may initiate mitral valve tissue (mal)adaptation (54). It is possible that the tricuspid valve, similarly, may (mal)adapt to repair. Thus, our and future work on tricuspid valve (mal)adaptation may also be used to optimize surgical repair techniques and interventional approaches to minimize post-operative (mal)adaptation.

## Limitations

Naturally, our study is subject to several limitations. Regarding our animal model, we chose a TIC biventricular heart failure model over a pulmonary hypertension model as it better represents the clinical setting of TR. That is, few patients present with isolated TR. In fact, 85-90% of severe TR cases are functional, resulting predominantly from left heart disease (1). Our model, with which we have years of experience, demonstrates all salient features of biventricular disease such as reduced left and right ventricular EF, dilated ventricles, mitral and tricuspid annular dilation, and regurgitation in both valves (24). However, our ovine model develops biventricular heart failure within weeks as opposed to years as in patients. Future studies should extend this time window. Also, the use of this model prohibits isolating the effects of mechanical stimulation (i.e., from annular dilation and chordal tethering) from hemodynamic stimulation. However, as our intent was to evaluate the valve’s propensity to (mal)adapt, in the presence of functional TR, we do not see this as a significant limitation. Nonetheless, future studies should isolate the underlying mechanisms through use of other models. Although limitations remain, they provide clear future directions for this line of research.

## Conclusions

In conclusion, we used an ovine TIC biventricular heart failure model to reveal that the anterior tricuspid valve leaflet (mal)adapted on multiple scales. Namely, we observed in the leaflets of these animals: i) an active metabolic protein response suggestive of increased energy expenditure, ii) TVAL cell expression of (mal)adaptive markers, iii) upregulated collagen synthesis, iv) reorganized collagen microstructure, v) increased tissue thickness, vi) increased leaflet area, and vii) increased stiffness. As in the mitral valve, we submit that these multi-scale changes are linked. Specifically, we borrow from our understanding of the mitral valve and propose that, as pathological stimuli activate valvular interstitial cells and lead to EndoMT, changes in cellular phenotype and cell proliferation likely emphasize inflammatory and profibrotic signaling pathways, which upregulate protein expression that initiates increased matrix turnover. As tissue-scale properties are dependent upon their micro-scale composition, the tissue consequently exhibits altered morphological and mechanical properties as a direct result of cellular scale activity.

## Abbreviations

αSMA: alpha smooth muscle actin
CTL: control
EF: ejection fraction
EndoMT: endothelial-to-mesenchymal transition
IDd: inner dimension at diastole
LV: left ventricle
MMP13: matrix metalloproteinase 13
RV: right ventricle
TGF-β1: transforming growth factor beta 1
TIC: tachycardia-induced cardiomyopathy
TR: tricuspid regurgitation
TVAL: tricuspid valve anterior leaflet
VIC: valvular interstitial cell

## Perspectives (Clinical Competencies/Translational Outlook)

### Clinical Competencies

*Medical Knowledge* - Valve extrinsic factors such as annular dilation and ventricular remodeling are well-established contributors to functional tricuspid regurgitation. Additionally, we suggest that tricuspid leaflet maladaptation be a potential novel disease mechanism for valvular incompetence, warranting further investigation.

### Translational Outlook

Future studies can identify the underlying mechanisms of leaflet maladaptation, its effect on valvular incompetence, and subsequent pharmaceutical therapeutics that aim to reduce its detrimental effects on valvular function.

## Funding Sources

Research reported in this publication was supported by the National Heart, Lung, And Blood Institute of the National Institutes of Health under Award Number F31HL145976 (WDM), the American Heart Association for their support under Award Number 18CDA34120028 (MKR), and internal grants from Meijer Heart and Vascular Institute at Spectrum Health. The content is solely the responsibility of the authors and does not necessarily represent the official views of the National Institutes of Health or the American Heart Association.

## Notes

### Competing Interest Statement

Marcin Malinowski and Tomasz Jazwiec are the Peter C and Pat Cook Endowed Research Fellows in Cardiothoracic Surgery from the Spectrum Health Foundation, Grand Rapids, MI. Manuel Rausch has a speaking agreement with Edwards Lifesciences, Irvine, CA. None of the other authors have any conflicts of interest to report.

