## Supplement for "The tricuspid valve also maladapts: A multiscale study in sheep with biventricular heart failure"

[‡]Division of Cardiothoracic Surgery, Spectrum Health, Grand Rapids, Michigan, USA.

[§]Department of Cardiac Surgery, Medical University of Silesia, School of Medicine in Katowice, Katowice, Poland.

[||]Department of Cardiac, Vascular and Endovascular Surgery and Transplantology, Medical University of Silesia in Katowice, Silesian Centre for Heart Diseases, Zabrze, Poland.

[¶]Department of Chemical Engineering, Texas Tech University, Lubbock, Texas, USA.

[#]Department of Aerospace Engineering and Engineering Mechanics, The University of Texas at Austin, Austin, Texas, USA

### Expanded Methods

#### Animal model, medications, and procedures

The tachycardia-induced cardiomyopathy (TIC) ovine model used in this study has been previously described in detail and validated as a reliable and repeatable model of biventricular heart failure in sheep (24). In short, we randomly assigned adult male Dorset sheep to either control (CTL, n = 17, 59.9 ± 4.6 kg) or disease (TIC, n = 33, 60.1 ± 5.3 kg) groups. Initially, all animals were acclimated for a 7-day period. In all animals, we monitored baseline (i.e. prior to pacing) ventricular function and valvular competence via epicardial echocardiography. We acquired all echocardiographic images with a 1.5 - 3.6 MHz transducer and Vivid S6 ultrasound machine (GE Healthcare, Chicago, IL). We used the American Society of Echocardiography criteria to assess valvular insufficiency. Using color flow and continuous wave Doppler, an experienced cardiologist graded TR as none (0), trace (0.5), mild (1.0), moderate (2.0), moderate-severe (3.0), or severe (4.0). We applied local anesthesia with 1% lidocaine for external right jugular intravenous (IV) catheter placement. We then initiated anesthesia (propofol 2-5 mg/kg IV), intubated, and mechanically ventilated each animal. We maintained general anesthesia via isoflurane inhalation (1%-2.5%) and fentanyl (5-20 µg/kg/minute). We monitored arterial blood pressure through an 18-gauge left carotid artery catheter (Teleflex, Morrisville, NC).

In TIC subjects, we sutured a monopolar pacing lead onto the lateral left ventricular wall, gaining access through a left lateral mini-thoracotomy approach (10-15 cm, fifth/sixth intercostal space) with a surgical incision approximated in standard fashion. Additionally, we infiltrated intercostal nerves with bupivacaine (0.25%). We exteriorized the lead through the thorax to a pacemaker (Consulta CRT-P, Medtronic, Minneapolis, MN). We stabilized the pacemaker in the subcutaneous pocket near the spine. Once the animal was breathing spontaneously, standing, and eating in the recovery area, we moved the animal to the pen. We provided prophylactic antibiotics preoperatively and postoperatively for 10 days (cefazolin 2 g IV q12H, and gentamicin 240 mg IV q24H). After 4-5 days of recovery, we initiated a progressive 180-260 beats per minute pacing protocol. During the pacing of TIC animals, we performed surveillance transthoracic echocardiography every three days to assess heart failure progression. One hour prior to each echocardiography study, we paused pacing, which was resumed at the conclusion of each study. Once both moderate tricuspid regurgitation (TR) and left ventricular dysfunction (ejection fraction < 30%) were present (19 ± 6 days), we performed a terminal epicardial echocardiograph.

During the terminal procedure of all animals, we acquired hemodynamic data over three full cardiac cycles via pressure transducers (PA4.5-X6; Konigsberg Instruments, Pasadena, CA) in the left and right ventricles via the apex, and the right atrium. We acquired cardiac pressure data and electrocardiographic recordings at 128 Hz over three cardiac cycles. We defined end-diastole as the time of the peak of the R-wave, and end-systole as the time of the maximum negative time derivative of left ventricular pressure. We euthanized all animals via sodium pentothal (100 mg/kg IV) and potassium chloride bolus (80 mEq IV). In CTL animals, the terminal procedure was performed after baseline epicardial echocardiograph. Lastly, we isolated the tricuspid valve from each animal.

For disclosure, we previously published the morphological, thickness, biaxial mechanics, and two-photon data for n = 6 CTL sheep included in this study (31). However, these data were all supplemented with additional subjects for this study, as the focus of that study was to report our findings in only CTL matched subjects. Additionally, to maximize utilization of each animal model, some TIC subjects were also used in a previously published study, unrelated to the leaflet (mal)adaptive response (25).

**Morphology and storage**

Upon TV isolation, we cut open the valve leaflet complex by separating the leaflets at the posterior-septal commissure to enable valve unfolding. We floated the TV, atrialis side up in 1XPBS and orthogonally photographed the TV leaflets on a calibrated grid. Using these photographs, an experienced cardiac surgeon identified the commissural points at which leaflets were separated and, using custom MATLAB code, we calculated the TVAL area, major cusp width, and major cusp height. All pixel measurements were translated to length metrics by means of the calibrated grid. We then cryogenically stored the tissue at -80°C in a 9:1 ratio of DMEM:DMSO with protease inhibitor until further testing. Once ready to be tested, we rapidly thawed the vials in room temperature water and removed all chordae tendineae from the ventricularis surface of the TV anterior leaflet (TVAL) prior to tissue sample collection for testing.

**High through-put analysis of protein expression**

We used TVAL fragments without consideration of localization/region for high-throughput mass spectrometry analyses. We finely minced the tissues and lysed cells at 4 $^{o}$C in a buffering solution containing 50 mmol/L HEPES (pH=7.4) supplemented with 150 mmol/L sodium chloride, 2 mmol/L dithiothreitol, 1% Igepal and a protease inhibitor cocktail (Active Motif, 100546). We incubated a total of six samples (3 CTL and 3 TIC) with mild agitation at 4 $^{o}$C for 2 hours. We determined the total protein concentration of each sample by bicinchoninic acid assay (Thermo, PI-23252). After quantification, we ethanol precipitated 50 µg of protein from each sample. After precipitation, we re-dissolved the proteins in Laemmli buffer for one-dimensional gel electrophoresis, Coomassie blue staining and in-gel digestion, following a previously published protocol (27). We dehydrated the gel pieces successively in 50% and 100% acetonitrile, and digested the proteins overnight in 0.1 µg sequencing-grade trypsin contained in 50 uL total volume. Finally, we extracted the peptides twice in 50% acetonitrile with 0.1% formic acid, which we dried to completion in a vacuum centrifuge.

We submitted the dried peptides to the University of Texas at Austin CBRS Biological Mass Spectrometry Facility for protein identification by liquid chromatography-tandem mass spectrometry (LC-MS/MS) using the Dionex Ultimate 3000 RSLCnano LC coupled to the Thermo Orbitrap Fusion (28). Prior to HPLC separation, we desalted the peptides using Millipore U-C18 ZipTip pipette tips following the manufacturer's protocol. We performed a C18 trap column, followed by a 75 µm I.D. x 25 cm long analytical column packed with C18 3 µm material (Thermo Acclaim PepMap 100) running a gradient from 5 - 35% acetonitrile. With a Fourier Transform Mass Spectrometry resolution of 120,000, and a 3 second cycle time, we acquired MS/MS in HCD ion trap mode. We uploaded the LC−MS/MS mzXML files MaxQuant (Max Planck Institute of Biochemistry, Germany) and searched against a protein database of Ovis aries downloaded from UniProt. We used default search parameters in MaxQuant for label-free quantification with a 1% false discovery rate. We selected only the proteins which had at least two replicates of intensity values across all treatment groups for the remaining analyses. With this, we confidently identified a total of 2457 proteins. We normalized intensity values per total protein content, and we set a fold-change of two prior to analysis in the Multi-Experiment Viewer (MeV, TM4 Microarray Software Suite). Using the non-parametric Wilcoxon Rank Sum statistical test to compare proteins in the two groups, we identified 247 significant differentially expressed proteins. We uploaded this list of proteins onto STRING v11 (Search Tool for the Retrieval of Interacting Genes/Proteins, STRING Consortium) for the generation of protein interaction maps. We further used the PANTHER (Protein ANalysis THrough Evolutionary Relationships, http://www.pantherdb.org/) classification system for the identification of Gene Ontologies (GO, http://www.geneontology.org). Refer to Supplementary Table 1 for all protein identifiers, protein family descriptions, and ontology classifications.

#### Histology, thickness, and immunohistochemistry

We fixed radial strips of TVALs from annulus to free edge in 10% Neutral Buffered Formalin for 24 hours and transferred them to 70% ethanol for storage until histological processing. We shipped the samples to a histological service (HistoServ, Inc., Amaranth, MD) for embedding, sectioning (5 um), and H&E staining. Using a light microscope (BX53 Upright Microscope, Olympus, Tokyo, Japan) with a 10x objective, we acquired and stitched full section images. Using custom MATLAB code, we fit splines to the atrialis and ventricularis surface of each section. We calculated the normal vectors along the atrialis spline, using these vectors to determine the distance between atrialis and ventricularis splines. Along the length, we summarized the thickness measurements into three regions (i.e. near-annulus, belly, and free edge), defined as three equidistant regions along the arc length of the atrialis spline.

Additionally, the commercial histological service (HistoServ, Inc., Amaranth, MD) performed five immunohistochemistry stains on the same radial strips. These stains include commonly used markers for growth and remodeling processes in previous studies (14,15,29,30): (i) α-smooth muscle actin (α-SMA) (Abcam, ab5694, Cambridge, MA, US) as a marker for valvular interstitial cell (VIC) activation, (ii) Ki-67 (Thermo Scientific, rb-1510-P0, Waltham, MA, US) to determine cell proliferation, (iii) matrix metalloproteinase 13 (MMP13) (Abcam, ab39012, Cambridge, MA, US) to quantify collagenases, and (iv) transforming growth factor β1 (TGF-β1) (Abcam, ab9758, Cambridge, MA, US) as a key profibrotic factor. Using the light microscope with a 40x objective, we acquired and stitched full section images. Using custom MATLAB code, we fit splines to the atrialis and ventricularis surface of each section. Next, we excluded regions of annular muscle, as annular muscle often skewed positive results. Along three equidistant length regions of the atrialis spline, again representing near-annulus, belly, and free edge regions, we interpolated 10 thickness regions between the atrialis and ventricularis splines - altogether splitting each radial strip section into 30 regions. We then passed each region of the image into a custom validated MATLAB program which detects positive and total pixels in that region, from which we determined the normalized percentage of positive stain in that region.

#### Quantitative collagen assay

We acquired the wet mass of tissue samples from annulus, belly, and free edge regions of TVALs. For every 10 mg of wet tissue mass, we added 100uL dH20 for homogenization. We hydrolyzed 100uL of homogenate in 100uL of 10N concentrated NaOH at 120°C for 1 hr, after which we neutralized by adding 100uL of 10N concentrated HCl. After vortex mixing at 2,000x g for 5 min, we transferred 10uL of hydrolysate to each well in triplicate, which we allowed to evaporate to dryness on a 65°C heating plate. We then followed the protocol provided with a total collagen assay kit (Biovision Inc., K406, Milpitas, CA, USA). From the quantitative assay, we measured the colorimetric absorbance at 560nm with a spectrophotometer (Tecan, Infinite 200 Pro,

Männedorf, Switzerland) which we interpolated from a collagen type I standard linear-fit curve ($R^2 = 0.99 \pm 0.01$). Note: Due to tissue shortage, this experiment was supplemented with 7 CTL sheep (male, Dorset, 46 ± 6 kg) TVALs which were not evaluated in animal model experiments. As collagen content is a normalized measure to tissue mass, we did not expect, nor acquire, differing data between these and other CTL animals.

**Biaxial testing and analysis**

We isolated 7mm x 7mm square samples from the belly region of TVALs, ensuring the major axes of the square aligned with the radial and circumferential directions of the leaflet. On the atrialis surface, we applied an approximate 3mm x 3mm grid of four ink fiducial markers in the tissue center to enable strain tracking during testing. To establish a stress-free reference configuration, we photographed these fiducial markers while the tissue floated in 1XPBS on a calibrated grid. We mounted each sample on a biaxial testing device (Biotester, Cellscale, Waterloo, ON, Canada) with rakes, ensuring the radial and circumferential axes of the tissue aligned with the axial directions of the device. After submerging the mounted samples in 37°C 1XPBS, we initiated a 10 cycle preconditioning force-controlled protocol to 300mN equibiaxially. After preconditioning we removed the tissue slack with a 10mN preload. We tested the tissue at 300 mN equibiaxially for two cycles each, during which we recorded (5 Hz) rake-to-rake distances, circumferential and radial forces, and images of the fiducial markers to compute local tissue stretches of the final downstroke. All biaxial tests were within four hours of tissue thawing.

Using the coordinates of the fiducial markers on the tissue throughout testing, we calculated the deformation gradient tensor, $\boldsymbol{F}$, with respect to the floating stress-free reference configuration. We then calculated the right Cauchy-Green deformation tensor, $\boldsymbol{C}$, via $\boldsymbol{C} = \boldsymbol{F}^T\boldsymbol{F}$, to acquire in-plane stretches throughout testing. Using the force data, we calculated the membrane tension as the force divided by the rake-to-rake distance in the orthogonal direction in the deformed configuration at each time point. In our analysis, we characterize these nonlinear J-shaped curves with four metrics: (i) toe stiffness, as the slope of the lower linear region of the curve, (ii) calf stiffness, as the slope of the upper linear region (near 20 N/m membrane tension) (iii) transition stretch, as the stretch at which the heel of the J-shaped curve is located (determined as the closest data point to the intersection of toe and calf stiffness lines), and (iv) anisotropy index, as the ratio between the circumferential and radial stretches at 20 N/m.

**Two-photon microscopy**

Post mechanical testing, we counterstained the 7mm x 7mm belly tissue samples for cell nuclei (Thermo Fischer Scientific, Hoechst 33342, Waltham, MA, US) for 20 minutes. Afterward, we optically cleared the tissue in an optical clearing solution (Glycerol:DMSO:5xPBS, 50:30:20%) under sonication for 30 minutes to improve imaging depth. We imaged three centrally located $500\mu$m x $500\mu$m regions through the entire tissue thickness at $10\mu$m z-steps with a two-photon microscope (Bruker, Ultima IV, Billerica, MA, US) and 20x water immersion objective (Olympus, XLUMPLFLN, Center Valley, PA, US). We imaged with a coverslip on the atrialis surface to avoid imposed stresses, and the tissue on a foil-lined glass slide to visually verify full-thickness image acquisition. We utilized second harmonic generation (SHG) at an excitation wavelength of 900nm and fluorescence at an excitation wavelength of 800nm to image collagen and cell nuclei, respectively. We epicollected the emission signal of collagen and cell nuclei with a photomultiplier tube channel filter (460 ± 25nm).

On each collagen image, we acquired coherency-based histograms of collagen fiber orientation by first normalizing the image histogram based on saturation and utilizing the ImageJ plugin OrientationJ on this processed image. Across the three z-stacks of each sample, we averaged and interpolated the OrientationJ output data into a single z-stack of normalized histograms from atrialis to ventricularis surfaces. From there, we fit von Mises distributions to each histogram acquiring the parameters $\mu$ and $\kappa$ representing the mean fiber angle and the fiber orientation concentration, respectively. Collectively, for every sample we imaged, we acquired a set of von Mises parameters representing mean fiber angles and fiber concentration at all depths. On each cell nuclei image, we acquired metrics of nuclear orientation, nuclear aspect ratio (NAR), and circularity by means of a custom MATLAB program which identifies individual nucleus contours in each image. With many nuclei in a single image, we acquire histograms of each of the three metrics at each depth of the z-stack, which we process in the same way as above. For nuclear orientation, we used von Mises distribution fits, while on NAR and circularity we fit normal distributions with parameters $\mu$ and $\sigma$ representing the mean and standard deviation, respectively.

#### Supplementary Figures and Figure Captions

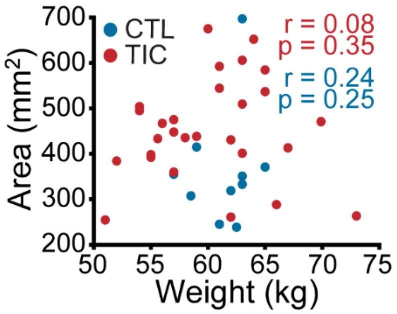


**<u>Supplementary Figure 1:</u> Tricuspid valve anterior leaflet area change does not correlate with animal weight.** Correlations between animal weight and anterior leaflet area with correlation coefficient, r, and p-value for tachycardia-induced cardiomyopathy (TIC, red, Pearson), and control (CTL, blue, Spearman) to ensure the changes we observed in leaflet area were not the result of animal weights.

(Figure file has additionally been separately attached online for size and legibility consideration, see file SupplementaryFigure2.tiff).

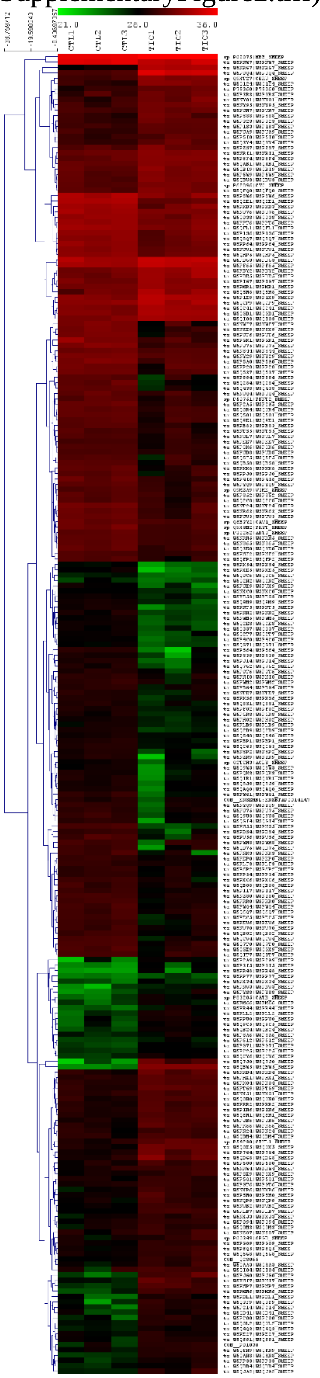


**<u>Supplementary Figure 2:</u> Proteomics revealed 247 differentially expressed proteins in tachycardia-induced cardiomyopathy (TIC) tricuspid anterior leaflets.** Heat map showing expression of significant ($p < 0.05$, fold change > 2) differentially expressed proteins between control (left 3 columns) and TIC (right 3 columns) groups. Each column represents a biological replicate within each group. Each row represents a protein, as marked by their uniport identifiers. Red, black and green correspond to high, medium and low intensity values, respectively. Proteins are clustered according to similarities in expression patterns and total intensities

(Figure file has additionally been separately attached online for size and legibility consideration, see file SupplementaryFigure3.svg).

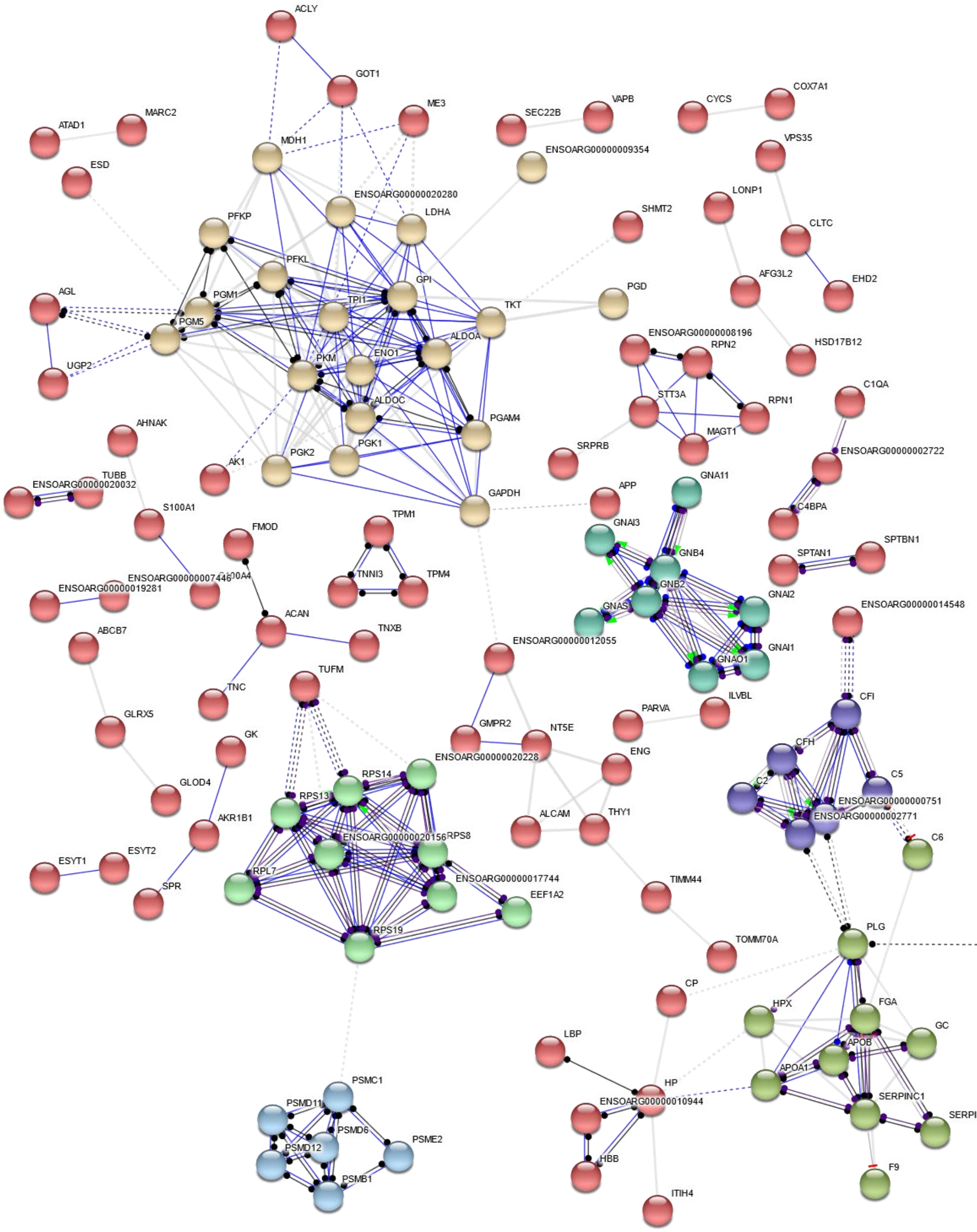


**Supplementary Figure 3: Interactome of 247 differentially expressed proteins showing clusters of shared action.** Proteins are interconnected on the basis of their molecular action upon each other. Clusters (marked with the same colors) were generated based on protein functional similarities. The nodes represent individual proteins while molecular action is represented by blue (binding) or black (reaction). Dashed associations represent low confidence interactions, grey associations indicate an undetermined molecular action. Distance between nodes are automatically generated based on functions. Disconnected nodes were hidden for simplicity. From top left, rotating clockwise, the clusters are metabolic, G proteins, complement proteins, serpins, proteasome and ribosomal proteins

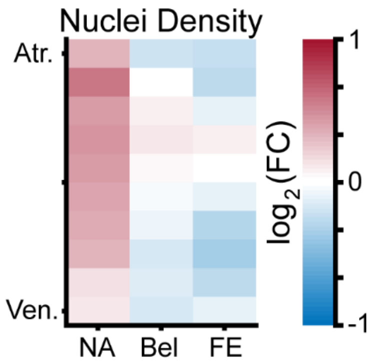


**<u>Supplementary Figure 4:</u> Marginal changes in TIC cellular nuclei density.** Heat maps showing regional changes in cell nuclei density. Heat map is separated by regions in radial (near-annulus (NA), belly (Bel), and free edge (FE)) and thickness (atrialis (Atr.), and ventricularis (Ven.)) axes. Fold change (FC) between control (CTL) and TIC was determined by the ratio of nuclei densities for each group. Color map indicates the logarithm base 2 of the FC, interpreted as: (positive, red) TIC nuclei density is higher than CTL, (0, white) TIC and CTL nuclei density are approximately equal, and (negative, blue) TIC nuclei density is less than CTL

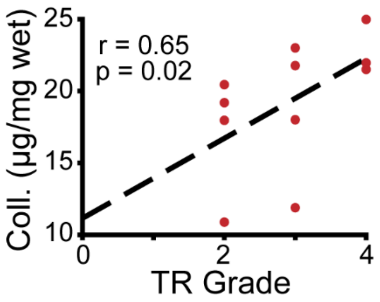


**Supplementary Figure 5: Collagen content positively correlates with clinical echocardiographic metric of tricuspid regurgitation (TR) severity.** Spearman correlation between echocardiography acquired TR grade with wet weight collagen content (Coll.) in tachycardia-induced cardiomyopathy (TIC) subjects. Correlation coefficients (r) and p-values are inscribed along with a linear fit (black, dashed) for visualization

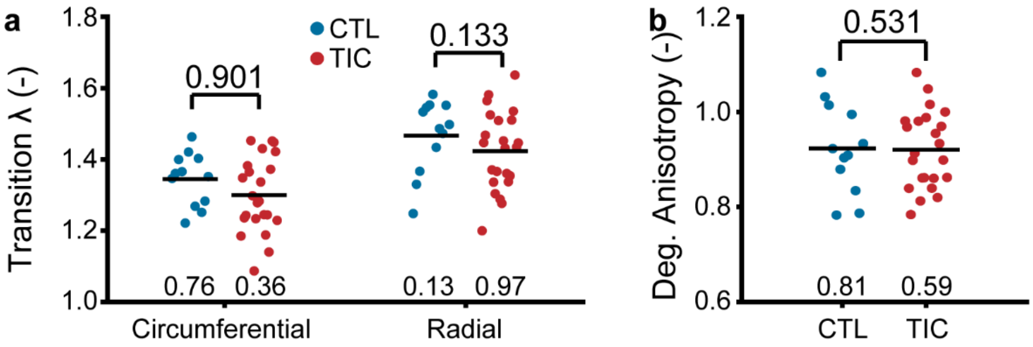


**<u>Supplementary Figure 6:</u> Biaxial curves between control (CTL, blue) and tachycardia-induced cardiomyopathy (TIC, red) exhibited comparible anisotropies. (A)** Comparisons of the transition stretch in circumferential and radial directions, defined as the stretch at which collagen engagement begins (i.e, the heel stretch). **(B)** Comparisons of the degree of anisotropy, defined as the ratio between circumferential and radial stretches at which 20 N/m membrane tensions are achieved under equibiaxial loading. Black bars represent data mean if normal, and data median if non-normal as detemined by Shapiro-Wilk test p-value below data. Values above data represent p-values from Student's t-test

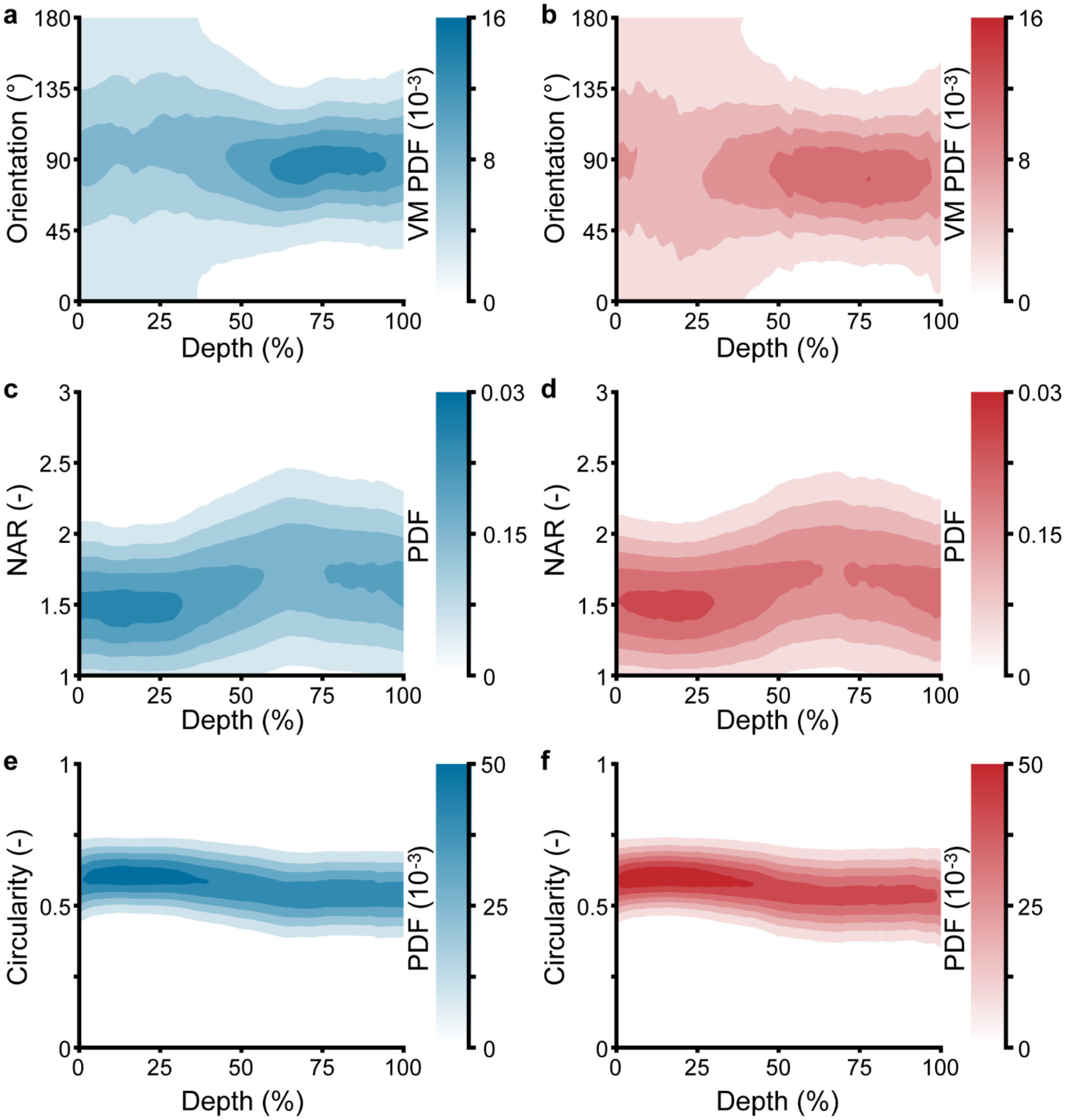


**<u>Supplementary Figure 7:</u> Through-depth nuclei microstructure reveals similar nuclei orientation, nuclear aspect ratio (NAR) and circularity throughout depth between control (blue) and tachycardia-induced cardiomyopathy (red). (A-F)** Heat map visualizations of von Mises probability distribution functions (VM PDF) and normal probability distribution functions (PDF) fit to the nuclei **(A, B)** orientation, **(C, D)** NAR, and **(E, F)** circularity histograms of two-photon acquired images throughout the entire depth (0% - Atrialis surface, 100% - Ventricularis surface) of averaged **(A, C, E)** control and **(B, D, F)** tachycardia-induced cardiomyopathy tissue samples. Orientations of 90° align circumferentially, while 0°/180° align radially. Nuclei microstructure remained qualitatively similar between groups with no significant findings

### Supplementary Tables

**<u>Supplementary Table 1:</u>** **(Sheet A)** List of FASTA headers, gene names, family information and expression levels for each of the 247 proteins. **(Sheet B)** Protein families identified, and their molecular functions, biological processes, cellular components, and protein class. **(Sheet C)** Chart summary of protein families identified and their common roles

(Table file has been separately attached online for size and legibility consideration, see file SupplementaryTable1.xlsx)
